# Loss of FOXP3 function causes expansion of two pools of autoreactive T cells in patients with IPEX syndrome

**DOI:** 10.1101/2022.07.10.499494

**Authors:** Šimon Borna, Esmond Lee, Uma Lakshmanan, Melissa Mavers, Mansi Narula, Akshaya Ramachandran, Jeanette Baker, Janika Schulze, Sven Olek, Louis Marois, Yael Gernez, Monica Bhatia, Alice Bertaina, Maria Grazia Roncarolo, Eric Meffre, Rosa Bacchetta

## Abstract

The monogenic autoimmune disease Immunedysregulation polyendocrynopathy entheropathy X-linked syndrome (IPEX) has elucidated the essential function of the transcription factor FOXP3 and of thymic-derived regulatory T (Treg) cells in controlling autoimmunity. However, the presence of autoreactive T cells in IPEX remains undetermined, thus representing a crucial gap in understanding the origin of autoimmunity in a FOXP3 deficient immune system. Combining epigenetic analysis as a lineage marker of Treg identity and TCR sequencing to assess the self-reactive clones, we showed that IPEX patients have two pools of expanded autoreactive T cells. The first originates from the expansion of autoreactive effector T cells (Teff), likely due to loss of Treg suppressive function since it is absent in carrier mothers, in whom Treg cells are functional. The second pool originates, unexpectedly, from Treg cells which lose their phenotypic markers, including CD25 and FOXP3. We call these loss of identity Treg cells and show that they are i) suppressed by healthy donor Treg in a patient post hematopoietic transplantation despite low donor chimerism, and ii) not detectable in patients with Autoimmune polyendocrinopathy-candidiasis-ectodermal dystrophy syndrome (APECED), a monogenic autoimmune disease of thymic origin. Moreover, we demonstrate that FOXP3 knock-out in Treg cells leads to increased Treg expansion and production of Th17 and Th2 cytokines, known to be increased in IPEX patients. These results suggest that the loss of identity Treg cells could directly contribute to immune dysregulation in IPEX. Collectively, we provide a better understanding of autoimmunity and novel ways to monitor the effects of Treg cell therapies in IPEX disease or other autoimmune diseases.

**One Sentence Summary:** Mutations of FOXP3 gene in humans cause expansion of autoreactive T cells originating from both effector T cells and regulatory T cells which gain effector function.

## INTRODUCTION

Autoimmune diseases affect more than 3% of people in western countries (*1, 2*) and their incidence is rising (*3, 4*). Autoimmunity occurs when immune cells fail to distinguish “self” from “non-self” and initiate immune reaction against their autologous tissues. The process of self-recognition is particularly important for T cells, because the specificity of the T cell receptor (TCR) is not fully encoded in the genome and TCR assembly gives rise to a broad range of TCR specificities, including autoreactive ones. To limit T-cell autoreactivity and thus prevent autoimmunity, developing T cells are selected in the thymus by medullary thymic epithelial cells (mTEC), B cells and dendritic cells, which express and present self-antigens to T cells (*5*). The expression of tissue restricted self-antigens in mTEC is regulated by autoimmune regulator (AIRE). Autoreactive T cells which recognize self-antigens with high affinity are eliminated (negative selection), while T cells which recognize the self-peptide/MHC complex with low affinity are positively selected and become effector T (Teff) cells (positive selection). Alternatively, some T cells with intermediate to high affinity to self-antigens may be positively selected and adopt a unique developmental fate by differentiating into regulatory T (Treg) cells. Hence, Treg TCR repertoire is physiologically self-reactive (*6–8*). However, the process of negative selection is not 100% accurate. Some autoreactive Teff cells escape negative selection and enter the periphery. Indeed, several reports demonstrated the presence of autoreactive T cells in human healthy donors (HD) (*9–11*). Consequently, a fraction of HD’s Teff cells proliferates in response to autoantigen stimulation *in vitro* and the addition of Treg cells inhibits their proliferation, demonstrating that Treg cells have a crucial role in controlling autoreactive T cell proliferation in humans (*9*).

Treg cell dysfunction was suggested to be one of the mechanisms responsible for the development of many autoimmune diseases (*12*). Two human monogenic diseases with autoimmunity well exemplify the importance of the thymic selection process and the consequences of the development of an aberrant immune tolerance. Mutations in AIRE lead to loss of expression of tissue-restricted antigens in thymus and result in failed T cell selection. Patients with mutations affecting AIRE function, develop Autoimmune polyendocrinopathy-candidiasis-ectodermal dystrophy syndrome (APECED), manifested by progressive autoimmune targeting of various tissues (*13*). Interestingly, analyses of Treg and Teff TCR repertoire of these patients indicated that some of the common Treg TCR clones were found lacking in the Treg compartment and present instead within the Teff compartment, indicating an altered distribution of the self-reactive T cells as a consequence of an aberrant selection (*14*). Similarly, mutations in FOXP3, a transcription factor essential for Treg function, lead to the prototypic example of Treg cell deficiency, called immune dysregulation polyendocrinopathy enteropathy X linked syndrome (IPEX), a severe early onset multiorgan life-threatening autoimmunity (*15*). However, T-cell autoreactivity in patients with IPEX has never been characterized, representing an important gap in understanding the role of FOXP3 and Treg cells in controlling T cell tolerance in humans.

In addition, growing evidences from murine models of autoimmune diseases and, to very limited extent, from human data, suggest that Treg cells that lose their stability due to a continuous exposure to an inflammatory environment may become Teff-like cells, alluding to but not demonstrating the existence of a second pool of peripheral pathogenic autoreactive T cells with different origin (*16–19*).

In contrast to the availability of lineage tracing in transgenic murine models, in humans it has been difficult to track Treg cells that have lost their phenotype and become Teff-like cells. Therefore, the stability of human Treg cells remains ill-defined. To overcome the traceability issue and to investigate the presence and origin of autoreactive T cells in IPEX, we used the epigenetic Treg marker named Treg specific demethylation region (TSDR) to track Treg cells independently of their current cell phenotype. Demethylation of TSDR is unique to FOXP3-expressing Treg cells and distinguishes them from Teff cells that can transiently express FOXP3 upon activation (*20*). Therefore, TSDR demethylation became the best marker for defining thymus-derived Treg cells. We have previously shown that patients with IPEX have an increased frequency of TSDR demethylated cells, suggesting that they have an expanded Treg cell compartment (*21*). In contrast, the frequency of Treg cells measured by their standard phenotypic markers CD3^+^CD4^+^CD25^high^ and FOXP3^+^ and/or CD127^-^ is normal to reduced (*22–25*). This unexpected discrepancy prompted us to hypothesize that a fraction of TSDR demethylated Treg cells lose their phenotypic markers, become Teff-like cells and are expanded in patients with IPEX.

Using TSDR demethylation as a Treg lineage marker, we show here that in contrast to HD, a fraction of IPEX Treg cells is detectable in the Teff compartment, demonstrating the presence of a population we named loss of Treg identity cells, since they no longer express the phenotypic markers of Tregs including CD25 and FOXP3. We further show that the loss of Treg identity is suppressed by healthy Treg cells in vivo, in studies of a patient post transplantation with partial donor Treg cell chimerism and in carrier mothers of FOXP3 mutations. Moreover, we combined the TSDR demethylation analysis with TCR receptor sequencing and showed, for the first time, that IPEX patients have increased autoreactivity in bona fide Teff cells, and that this increase in autoreactivity has peripheral origin. Notably, the presence of loss of Treg identity cells strongly correlates with TCR autoreactivity of Teff cells, suggesting that loss of Treg identity cells may play a role in disease pathology. Indeed, using CRISPR/Cas9 mediated FOXP3 knock out in Treg cells, we demonstrate that loss of FOXP3 expression leads to increased Treg expansion and production of proinflammatory, Th2 and Th17 polarizing cytokines, recapitulating the IPEX patient immune dysregulation. Collectively, we show that loss of FOXP3 function in humans leads to increased autoreactivity in Teff cells and expansion of Treg cells which loss their identity, demonstrating expansion of two pools of autoreactive T cells in IPEX patients.

## RESULTS

This study was performed on samples from patients with IPEX syndrome, carrier mothers, and patients with APECED syndrome. Mutations, age, and sex of subjects included in the study are summarized in Table 1.

**Table 1.**
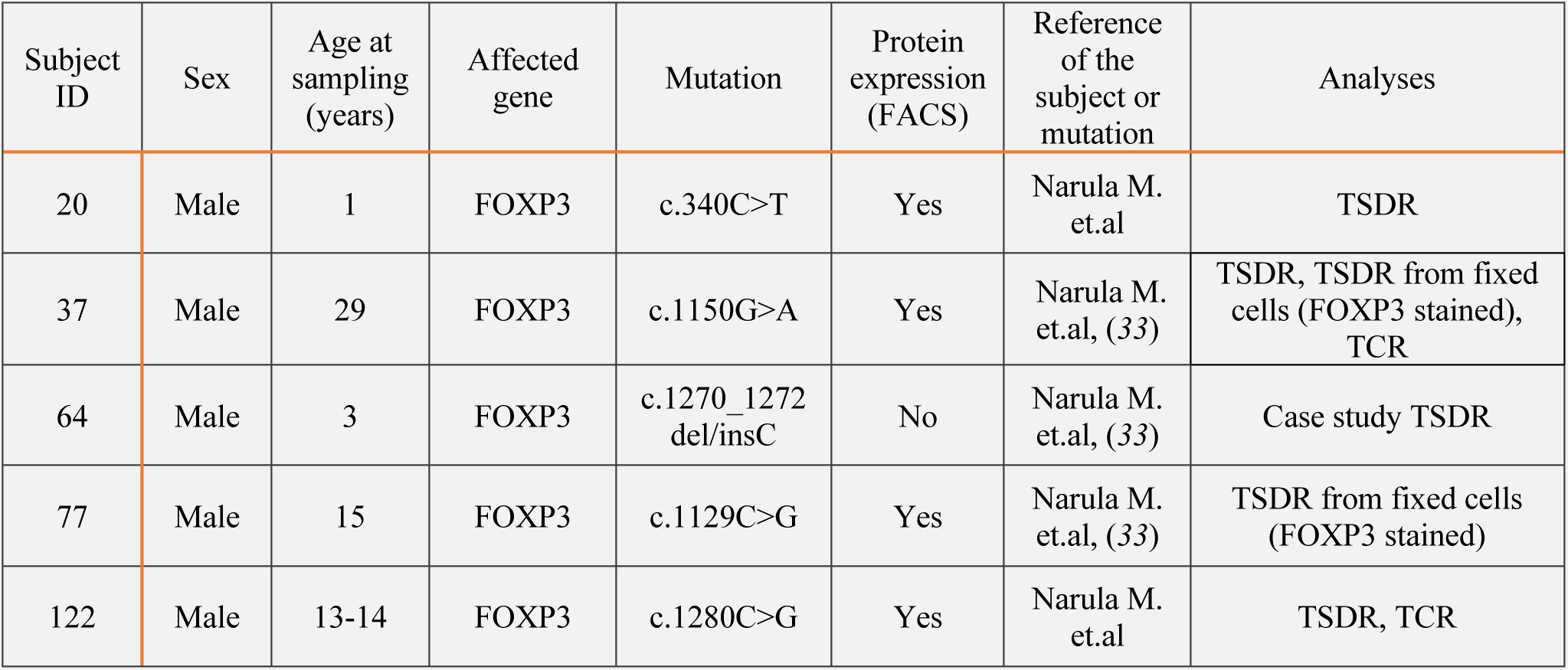
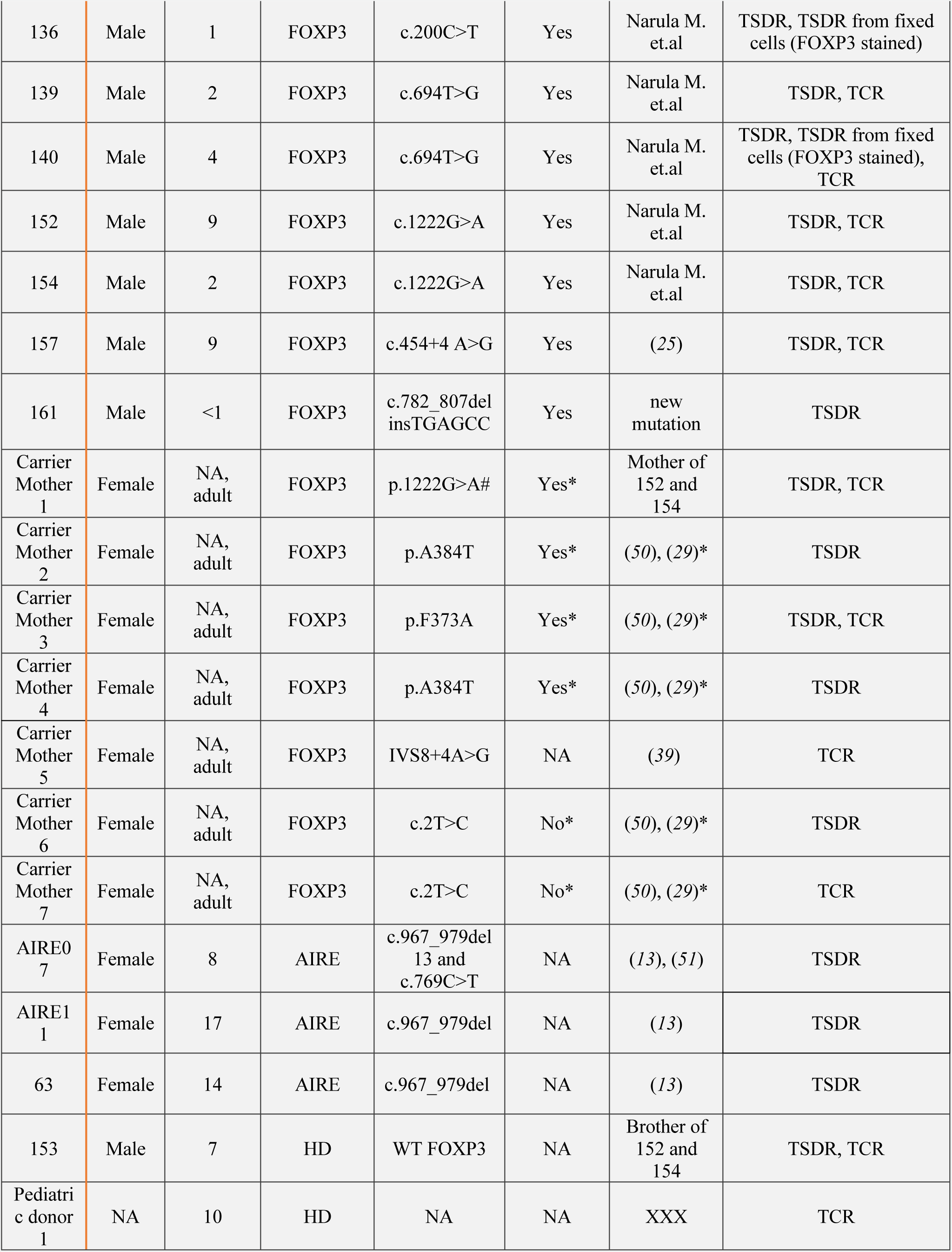
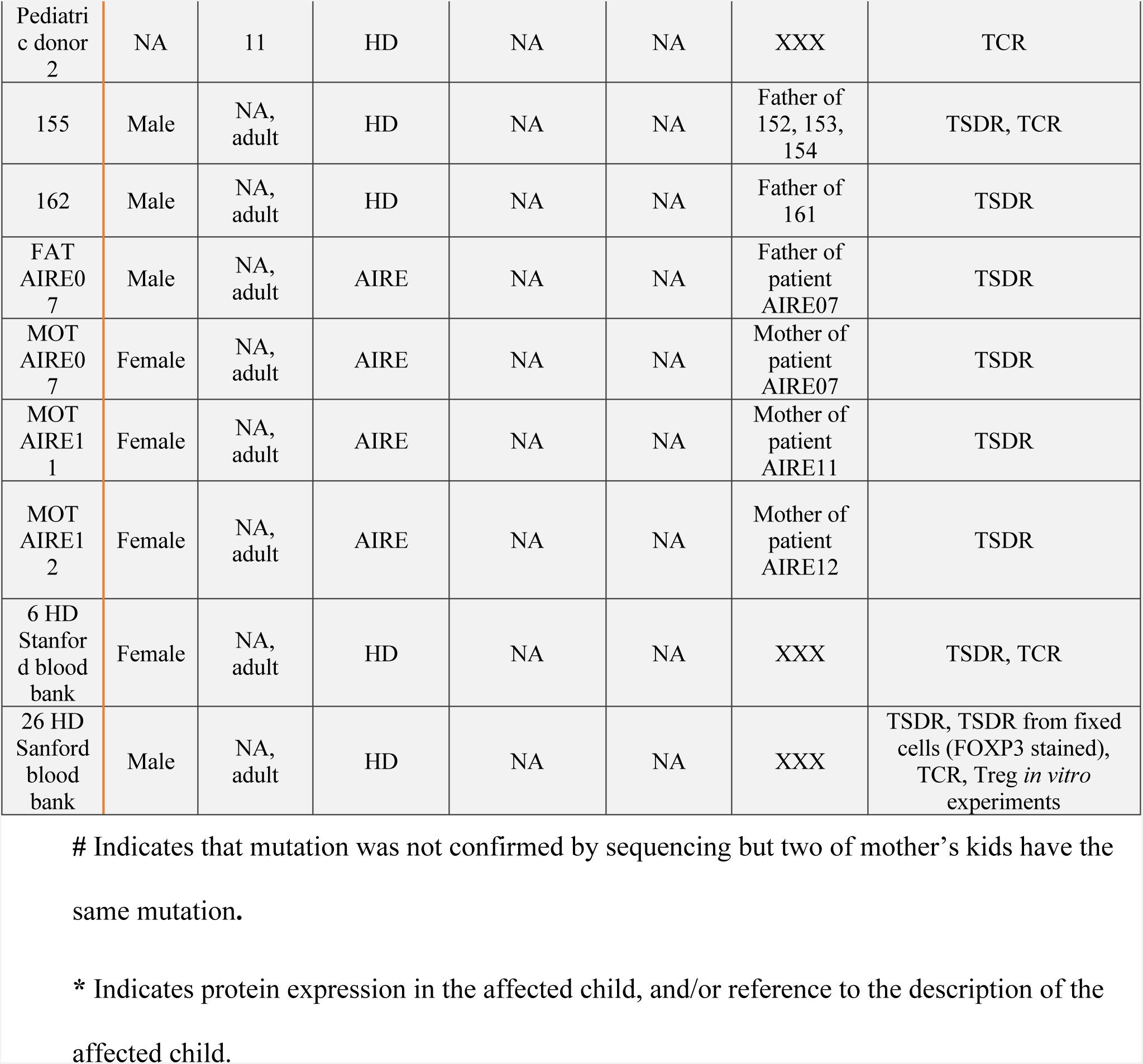
List of subjects analyzed in this study.

### Subhead 1: Mutations in FOXP3 lead to loss of Treg identity in vivo

To investigate the origin of the high frequency of TSDR demethylated CD4^+^ T cells in the whole blood of IPEX patients and understand the discrepancy between this finding and the low/normal frequency of CD3^+^CD4^+^CD25^high^CD127^low^ Treg cells detected by flow cytometry, we measured the TSDR demethylation in sorted CD3^+^CD4^+^ T cell subsets with different expression of CD25 and CD127, the relevant receptors for IL-2 and IL-7 cytokines differentially regulating Treg and Teff homeostasis (*26*). To this aim, two subpopulations of Teff were sorted as CD25^-^CD127^low^ and CD25^-/dim^CD127^high^, hereafter named Teff1 and Teff2, respectively (Fig. 1A). In parallel CD25^high^CD127^low^ Treg cells were isolated and shortly expanded in vitro to obtain sufficient amount of DNA for the TSDR demethylation analyses. The frequencies of Treg, Teff1, and Teff2 populations were not significantly different between HD and IPEX (Fig. 1B), and we did not observe expansion of CD4^+^ T cells in IPEX patients (Fig. 1C). Quantification of the TSDR demethylation showed that the sorted CD25^high^CD127^low^ Treg subsets of both IPEX and HD were fully demethylated whereas, in contrast to HD, a significant fraction of IPEX Teff1 cells were TSDR demethylated. This result demonstrates that a pool of cells with the FOXP3 epigenetic marker of Treg cells was comprised in the CD25 negative fraction, suggesting that a fraction of Treg cells had lost CD25 expression and are present among Teff1 cells (Fig. 1D). Moreover, although not reaching level of statistical significance between the groups of IPEX and HD, in some patients we could detect the presence of TSDR demethylated cells also in Teff2 population, suggesting that some of the TSDR demethylated cells could also gain CD127 expression, and therefore likely responding to different homeostatic regulation.

**Fig. 1.**
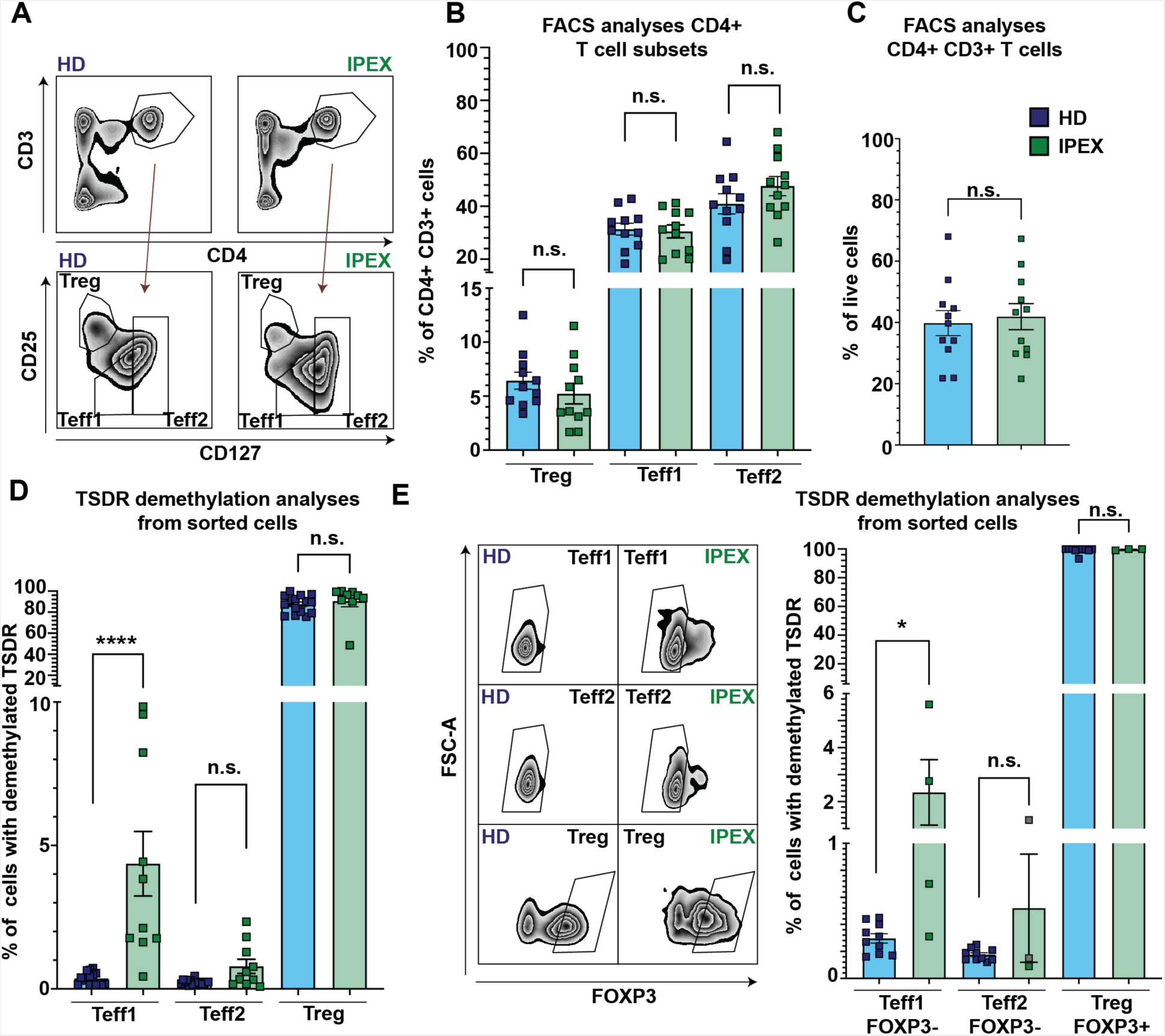
A fraction of Treg in IPEX patients loses Treg phenotypic markers. **(A)** Representative image of gating strategy used to sort and analyze frequencies of Treg, Teff1 and Teff2 subpopulations. **(B,C)** Quantification of frequencies of Treg, Teff1, Teff2, and CD3^+^CD4^+^ T cells from PBMC of 11 IPEX patients and 11 HD. **(D)** TSDR demethylation analyses from sorted Treg, Teff1, and Teff2 populations of 10 IPEX patients and 16 HD (sorted as shown in **A**). **(E)** TSDR demethylation analyses of FOXP3-Teff1 and Teff2 populations and FOXP3+ Treg from 4 IPEX patients and 10 HD. (*Left panel*) Representative image of gating strategy for FOXP3, of Teff1, Teff2, and Treg cells gated as shown in A. (*Right panel*) Quantification of TSDR demethylation. Data points shown in grey (IPEX Teff2) represents samples with low cell number/genome copies, where the values may be partially imprecise. Significance was evaluated using Mann Whitney test.

Since the vast majority of IPEX patients’ mutations do not abolish FOXP3 expression but diminish FOXP3 function, we tested if the loss of CD25 expression (and/or gain of CD127 expression) in IPEX Treg is also associated with loss of FOXP3 expression. We sorted FOXP3^+^ Treg and FOXP3^-^ Teff1 and FOXP3^-^ Teff2 cells and analyzed TSDR demethylation (IPEX: n=4 and HD: n=10) (Fig. 1E, S1). Remarkably, we found TSDR demethylated cells among FOXP3^-^ Teff1 in IPEX patients but not in HD. We were not able to obtain sufficient TSDR demethylation data, which would pass our internal quality control, to evaluate presence of TSDR demethylated cells in Teff2 population due to combination of low cell count and level of TSDR demethylation. Collectively, our data show that in IPEX patients a fraction of epigenetically determined Treg cells do not express CD25 and FOXP3, and some gain CD127 expression. These data explain the discrepancy between the frequencies of Treg measured by TSDR demethylation and FACS analyses using CD25 and CD127 markers in these patients and show that FOXP3 mutated Treg cells lose their phenotypic identity, demonstrating the presence of loss of identity Treg cells in IPEX disease.

### Subhead 2: IPEX patients have increased TCR repertoire autoreactivity and differential length usage of CDR3B

We determined the TCR repertoire in CD4^+^ T cells from IPEX patients to potentially illustrate in humans the FOXP3-dependent role of Treg in controlling autoreactive T cell expansion in vivo. We performed DNA sequencing of TCR complementary determining region 3 beta (CDR3B) of Treg, Teff1 and Teff2 cells from seven IPEX patients and ten HD. In total, we identified 5.4×10^5^ and 7.4×10^5^ Teff1, 5.1 x10^5^ and 6.9×10^5^ Teff2, 1.7 x10^5^ and 2.3 x10^5^ Treg TCR sequences from IPEX patients and HD, respectively. The number of sequences identified per sample for each subpopulation was well balanced between the IPEX and HD samples (Fig. 2A). The fraction of productive rearrangement from the total number of sequences identified was slightly reduced in all three populations of IPEX patients as compared to those of HD, although the difference was not statistically significant (Fig. 2B). In IPEX, the overall Simpson clonality in all three major populations was normal (Fig. 2C), but a trend toward higher clonality in IPEX Treg cells were observed in some patients. In addition, we did not find any major bias in TCRB gene usage (Fig. S2). However, alteration of TCR repertoires of Teff1 and especially Teff2 from IPEX patients were evidenced by their different distribution of CDR3 length compared to counterparts in HD. Indeed, while Treg cells from both IPEX patients and HD display similar CDR3 length distribution, Teff1 and Teff2 cells in IPEX were enriched in clones with longer CDR3B loops (Fig. 2D). To investigate TCR autoreactivity, we analyzed the presence of autoreactivity promoting amino acid (AA) doublets at positions 6 and 7 of the CDR3B loop as previously described (*7, 27, 28*). We found increased frequencies of the autoreactivity promoting AA doublets in the TCR repertoires of both Teff1 and Teff2 cells from IPEX patients (Fig. 2E). Consistently with the increased frequency of autoreactivity promoting AA doublets in IPEX Teff2 cells, we found significantly reduced frequency of autoreactivity limiting AA doublets in these cells and a trend toward reduced frequency in the Teff1 compartment, which, however, didn’t reach statistical significance (Fig. 2F). We conclude that differential length usage of CDR3B, increased frequency of autoreactivity AA promoting doublets, and reduced frequency of autoreactivity limiting AA doublets in Teff cells from patients with FOXP3 mutation reveal increased autoreactivity in the Teff compartments in IPEX. While Teff1 compartment is composed of both TSDR demethylated cells and bona fide TSDR methylated Teff, Teff 2 compartment contains only a negligible amount of TSDR demethylated cells. Therefore, we conclude that TCR repertoire autoreactivity in Teff2 compartment is originating from TSDR methylated bona fide Teff cells.

**Fig. 2.**
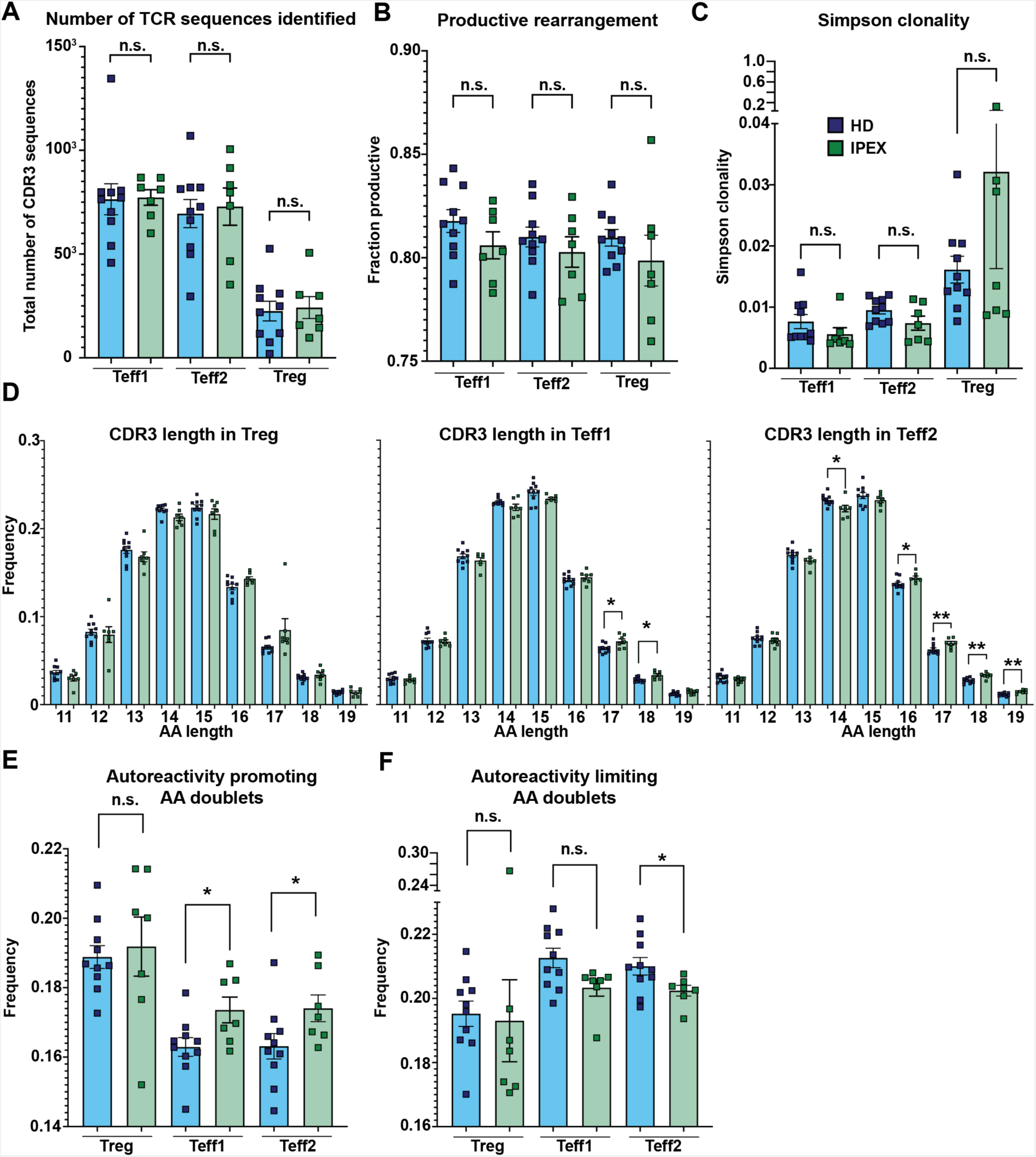
IPEX patients have increased TCR repertoire autoreactivity and differential length usage of CDR3B. **(A)** Similar number of TCR CDR3B sequences identified between IPEX patients and HD in Teff1, Teff2 and Treg populations. **(B)** Normal, or slightly but not significantly reduced fraction of TCR*β* productive rearrangement between IPEX patients and HD in Teff1, Teff2 and Treg populations. **(C)** Similar Simpson clonality between IPEX patients and HD in Teff1, Teff2 and Treg populations. **(D)** Analyses of CDR3B length of Treg (left), Teff1 (middle) and Teff2 (right) from IPEX patients and HD. **(E)** Frequency of autoreactivity promoting doublets and **(F)** autoreactivity limiting doublets of TCR CDR3B in IPEX patients and HD in Teff1, Teff2 and Treg populations. TCR CDR3B sequencing was performed in all three T cell subsets of 10 HD and 7 IPEX patients (A-F). Analyses in C-G were performed from productive rearrangements. Statistical analyses were performed using Mann Whitney test.

### Subhead 3: Frequency of autoreactive AA doublets in Teff cells positively correlates with loss of Treg identity in IPEX patients

To address whether the expanded pool of TSDR demethylated cells correlates with the increased TCR autoreactivity in IPEX patients, we correlated in Teff1 compartment, the presence of autoreactivity promoting AA doublets and the frequency of TSDR demethylated cells. We found a positive correlation between TSDR demethylation and TCR autoreactivity (p=0.0048), which could be explained by the presence of *loss of identity Treg cells* within that compartment (Fig. 3A).

**Fig. 3.**
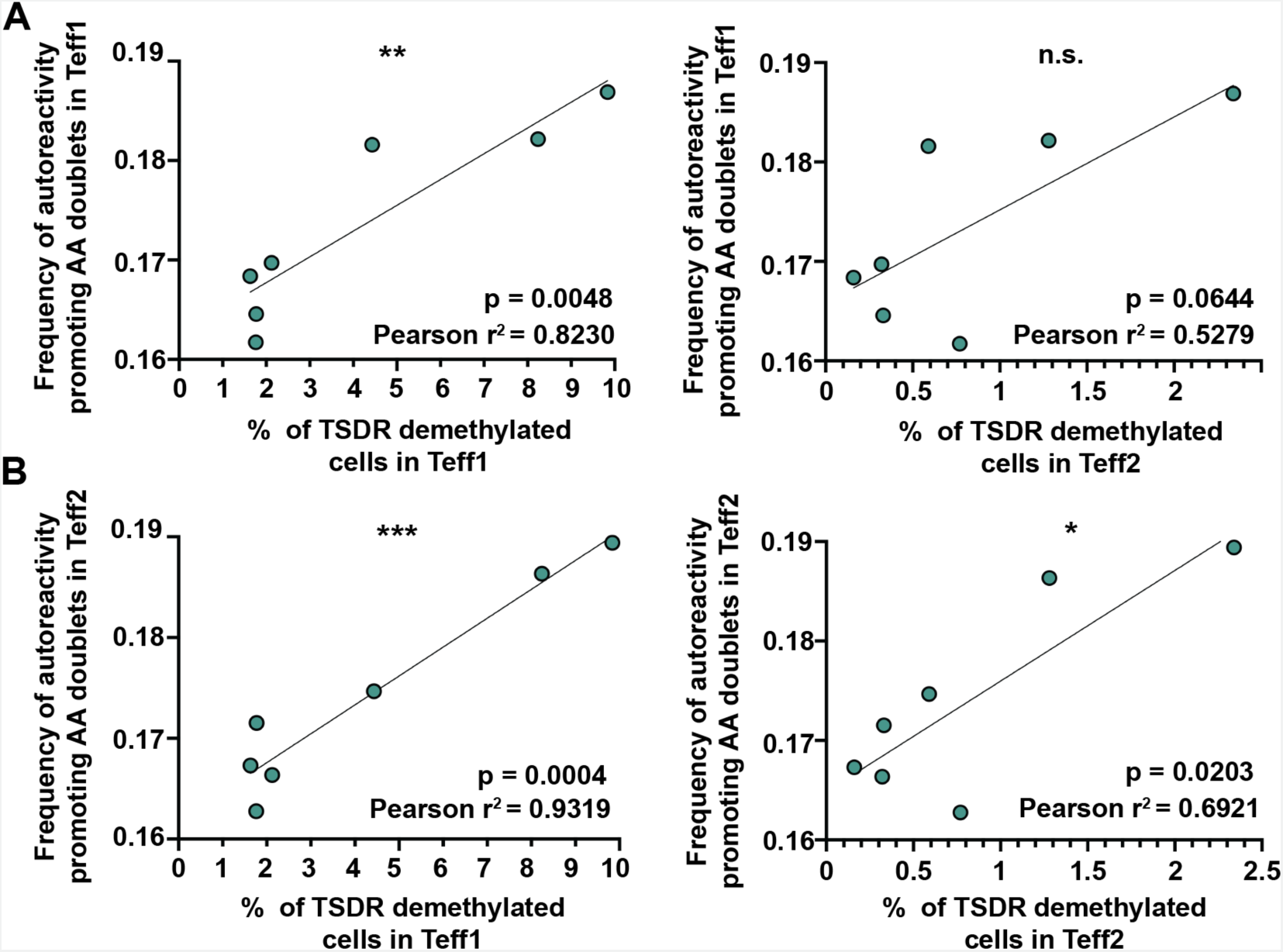
Frequency of TSDR demethylated cells in Teff compartments correlates with increased TCR receptor autoreactivity. **(A)** Correlation between % of TSDR demethylated cells in Teff1 (left) and Teff2 (right), and frequency of autoreactive doublets in Teff1. **(B)** Correlation between % of TSDR demethylated cells in Teff1 (left) and Teff2 (right), and frequency of autoreactive doublets in Teff2. Data were analyzed using Pearson correlation, and p value, Pearson r^2^, and a simple linear regression are shown (n=7).

Correlation between presence of autoreactivity promoting AA doublets in Teff2 and TSDR demethylation in Teff2 was also significant but weaker than in Teff1 (p=0.0203, Fig. 3B). These data are in line with very low numbers of TSDR demethylated cells in Teff2 compartment.

In addition, we found strong correlation between presence of TSDR demethylated cells in Teff1 and the frequency of autoreactivity promoting AA doublets in Teff2 compartment (p=0.0004). These data strongly suggest that the presence of TSDR demethylated cells in Teff1 compartment is predictive of increased autoreactivity of the Teff2 compartment composed in vast majority of bona fide Teff cells. Hence, loss of Treg identity may favor the expansion of autoreactive Teff cells.

### Subhead 4: Mothers of IPEX patients carrying *FOXP3* mutation do not show loss of Treg identity and display normal TCR autoreactivity

The *FOXP3* gene is located in the X chromosome. As a consequence, in carrier mothers of FOXP3 mutation, random chromosome X inactivation results in the expression of mutated FOXP3 in 50% of Teff cells (*29, 30*). In contrast to Teff, the majority of carrier mothers’ CD25^high^ Treg cells express the non-mutated *FOXP3* allele (*29, 31*). Therefore, in the mothers unlike in IPEX patients, the Treg compartment is functional and able to control the partially mutated Teff compartment. However, it is unknown i) if some FOXP3-mutated former Treg cells might be present in the Teff compartment as the loss of Treg identity, and ii) if carrier mothers have increased autoreactivity in the Teff compartment. First, similarly to IPEX patients, we sorted Treg, Teff1 and Teff2 cells from five carrier mothers, five sex-matched HD, and analyzed TSDR demethylation. We did not find TSDR demethylated cells in any of the Teff cell subsets tested (Fig. 4A). These results demonstrate that in carrier mothers, in which functional Treg cells are present, the loss of Treg identity cells are not detectable.

**Fig. 4.**
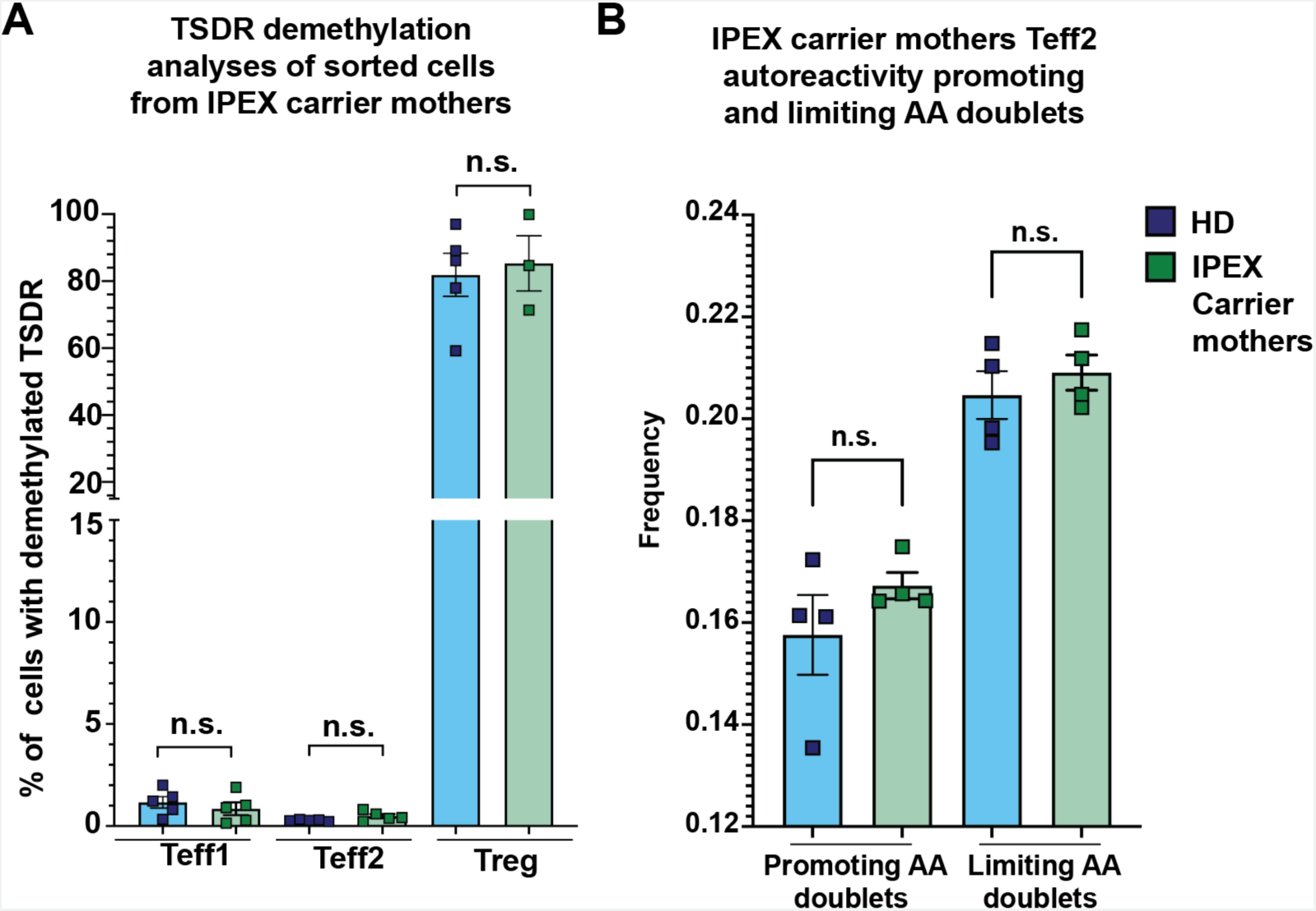
IPEX carrier mothers have normal TCR repertoire autoreactivity and distribution of TSDR demethylated cells in Treg and Teff compartments. **(A)** TSDR demethylation analyses from sorted Teff1, Teff2, and Treg populations from 5 FOXP3 mutation carrier mothers and 5 sex matched HD. Sorting strategy was the same as in figure 1A. **(B)** Frequency of autoreactivity promoting and limiting doublets in TCR CDR3B repertoire of Teff2 cells from 4 FOXP3 carrier mothers and 4 sex matched HD. Analysis in B was performed from productive rearrangements. Significant was evaluated using Mann Whitney test.

Then we investigated the frequency of T cell autoreactivity in the mothers to address, whether the increase in Teff autoreactivity in IPEX patients results from Teff extrinsic effect mediated by loss of Treg function or rather from a FOXP3 intrinsic role in Teff development (*32*). We sequenced TCR CDR3B repertoire of Teff2 from four carrier mothers and four sex-matched HD (Fig. 4B). Although only half of the carrier mothers Teff cells carry FOXP3 mutation, we decided to perform bulk TCR repertoire profiling to obtain a sufficient number of sequences for this analysis. We assumed that if the increased frequency of autoreactivity promoting AA doublets stems from aberrant T cell development and may be present in 50 % of carrier mother’s T cells, we may be able to observe a trend toward increased and reduced frequency of autoreactivity promoting and limiting doublets, respectively. We did not find any differences, nor a trend, in frequencies of autoreactivity promoting and limiting AA doublets in carrier mothers, as compared to healthy donors, supporting that the increased TCR autoreactivity in IPEX Teff cells mainly originate from the peripheral expansion of autoreactive TCR clones owing to a loss of Treg function.

### Subhead 5: Low frequency of loss of identity Treg cells in Teff and absence of increased frequency of TSDR demethylated cells in a FOXP3 null IPEX patient after haemopoietic stem cell transplantation (HSCT) despite low donor chimerism

Since loss of Treg identity is present only in patients with IPEX disease, but not in carrier mothers, we tested if the presence of loss of Treg identity cells is suppressed by functional Treg cells *in vivo* in a IPEX patient with mixed donor chimerism, post allogeneic HSCT. We analyzed frequency of TSDR demethylated cells in the Teff compartment of a FOXP3 null IPEX patient prior to and 2.5 years post HSCT. The patient’s and the donor’s cells could be distinguished using FACS by the mismatched expression of HLA-A2. The patient donor chimerism 2.5 years post transplantation was only about 15% in Teff cells and about 38% in Treg cells (Fig. 5A). Despite the relatively low chimerism at this time post-HSCT in both Treg and Teff cells, we observed a three-fold reduction in frequency of TSDR demethylated cells in the host Teff compartment as compared to the pre-HSCT sample (Fig. 5B). In addition, the kinetics of the TSDR demethylation in whole blood showed that the proportion of TSDR demethylated cells relative to frequencies of CD4^+^ T cells significantly decreased overtime reaching normal values, one year post HSCT (Fig. 5C, D). These data suggest that presence of even a small proportion (about 40%) of Treg cells of donor origin prevents the expansion of IPEX Treg cells following loss of Treg identity.

**Fig. 5.**
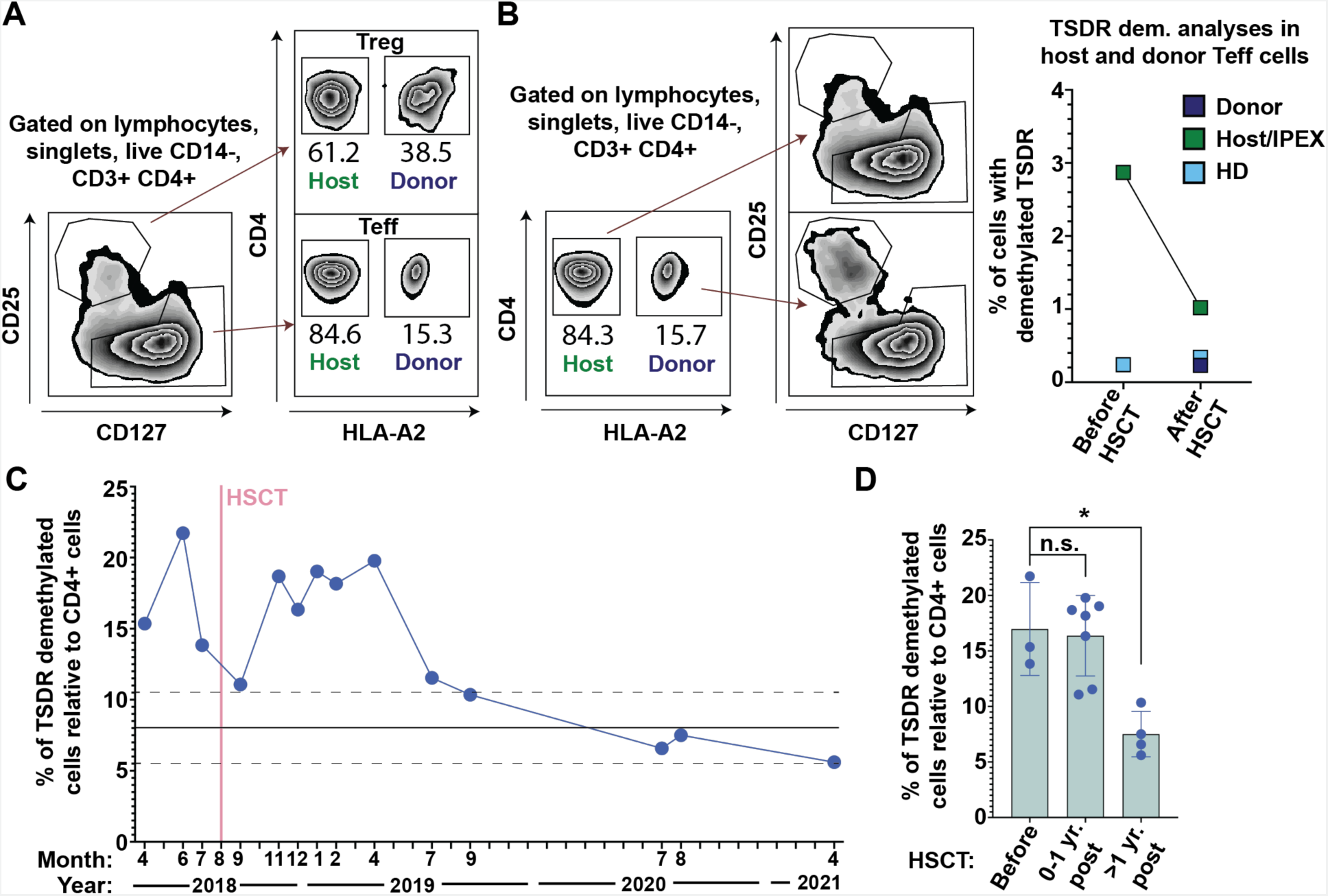
Presence of HD graft prevents expansion of TSDR demethylated cells and loss of Treg identity in an IPEX patient post haemopoietic stem cell transplantation despite low donor chimerism. **(A)** FACS analysis of IPEX patient chimerism 2.5 years post transplantation show competitive advantage of healthy donor Treg cells over the patient’s Treg. Donor cells are HLA-A2^+^ and host cells are HLA-A2^-^. **(B)** TSDR demethylation analyses of sorted host and donor Teff from patient before and 2.5 years post transplantation. In addition, HD Teff cells were sorted in parallel as a control (light blue). (*Left panel*) Gating strategy. (*Right panel*) Results from TSDR demethylation analyses. **(C)** TSDR demethylation analyses from whole blood at various time points pre- and post-transplantation. TSDR demethylation was normalized to % of CD4^+^ T cells also determined by epigenetic analyses. Mean and standard deviation of HD (internal data of Bacchetta lab) are depicted as full line and dashed lines, respectively. **(D)** Quantification of TSDR/CD4 ratio before, less than 1 year post transplantation, and more than one year post transplantation. Data were analyzed using Dunn’s multiple comparisons test.

### Subhead 6: Loss of Treg identity of IPEX Treg is not present in APECED patients

We speculated that in the absence of functional FOXP3, TSDR demethylated Treg precursors fail to commit to the Treg lineage during mTEC-mediated thymic selection and become instead TSDR demethylated Teff-like cells. To get insight into the potential thymic origin of loss of Treg identity, we analyzed TSDR demethylation in Treg and Teff cells from APECED patients, in which a T cell extrinsic defect in thymic selection leads to impaired Treg commitment and presence of clones that were meant to be Treg, in the Teff compartment (*14*). We sorted Treg, Teff1 and Teff2 cells from three APECED patients and five HD or carrier relatives if available. In contrast to IPEX patients, TSDR demethylated cells were not detected in the Teff compartment of APECED patients, whereas APECED’s Treg cells were fully TSDR demethylated (Fig. 6A). In line with the aberrant Treg and Teff cells selection, analyses of previously reported TCR sequencing data from four APECED patients and four HD (*14*) showed increased frequency of autoreactivity promoting AA doublets in both Treg and Teff cells of APECED patients as compared to the HD. However, in Teff cells the data were borderline line of statistical significance (Fig. 6B, p-value = 0.0571). The frequency of autoreactivity limiting AA doublets in Teff cells were also reduced and showed the same borderline of statistical significance. Hence, although APECED patients present with increased Teff cells’ TCR autoreactivity, the Treg cell lineage commitment in AIRE-deficiency does not result in the aberrant presence of TSDR demethylated cells in the peripheral Teff compartment of APECED patients. These results therefore suggest that TSDR demethylated Teff cells are specifically induced by defective FOXP3 function likely through peripheral loss of Treg cell identity.

**Fig. 6.**
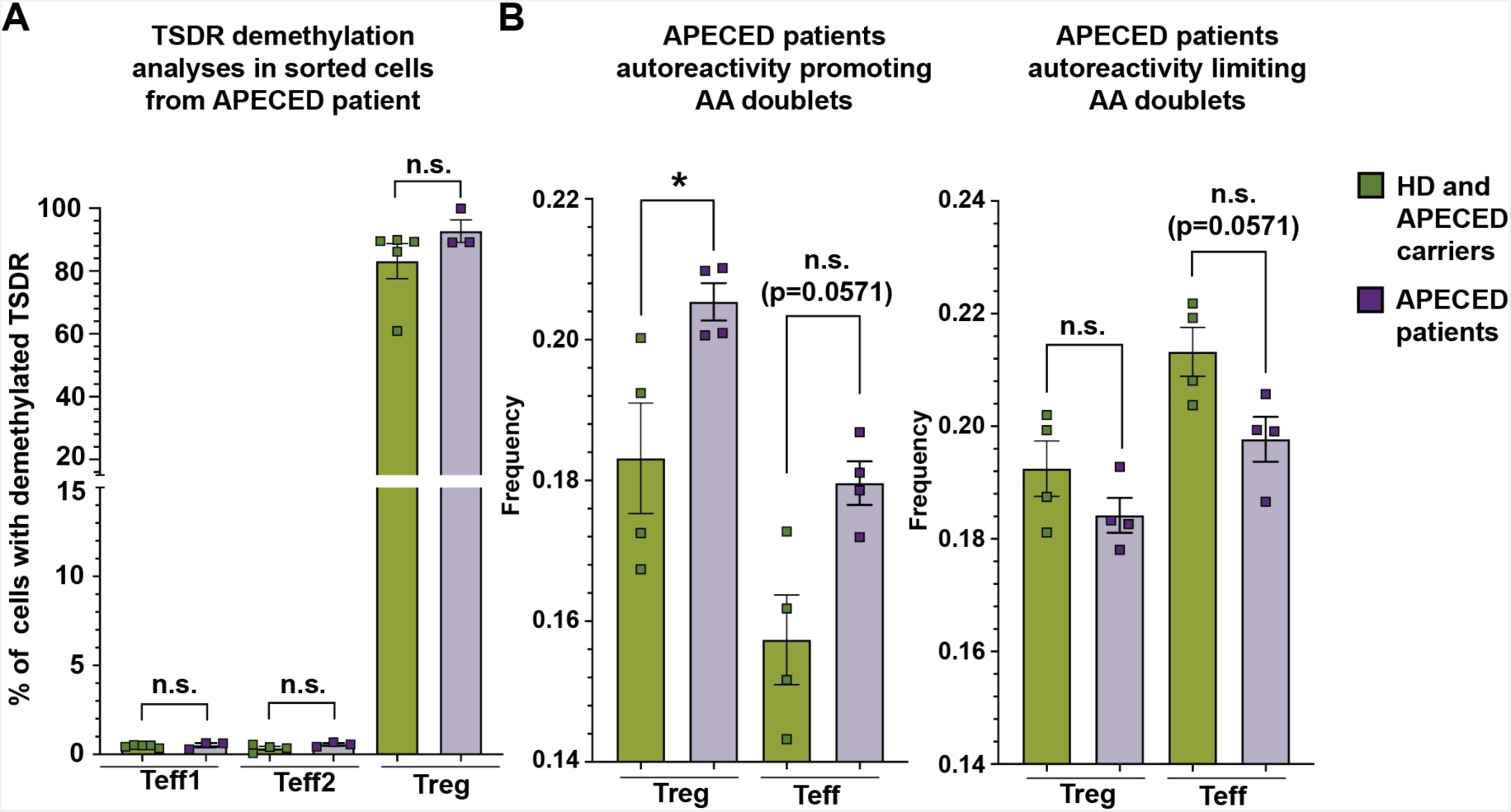
APECED patients have increased TCR repertoire autoreactivity but a normal distribution of TSDR demethylated cells in Treg and Teff compartments. **(A)** TSDR demethylation analyses from sorted Teff1, Teff2 and Treg populations from 3 APECED patients and 5 healthy controls including 4 parents and 1 HD. **(B)** Analyses of TCR receptor autoreactivity from previously reported TCR sequencing data of four APECED patients and four healthy donors (*14*). (*Left graph*) Frequency of autoreactivity promoting AA doublets in TCR CDR3B of Treg and Teff cells. (*Right graph*) Frequency of autoreactivity limiting AA doublets in TCR CDR3B of Treg and Teff cells. P value for borderline significant data is indicated. Analyses was done from productive rearrangements. Data were analyzed using Mann Whitney test.

### Subhead 7: Greater expansion and increased production of proinflammatory cytokines in FOXP3-knocked out Treg in response to TCR stimulation in vitro

IPEX patients have increased frequencies of TSDR demethylated cells and a fraction of these Treg have lost Treg phenotypic markers including FOXP3 expression. We investigated the Teff function of Treg following loss of FOXP3 expression. To overcome the limited availability of IPEX patients’ Treg cells, we prepared FOXP3 knock out (KO) Treg from HD Treg using our previously described knock in - knock out strategy (*33*). Briefly, we used CRISPR/Cas9 technology combined with AAV6 delivery of FOXP3 homologous template containing NGFR reporter gene under control of PGK promoter. We isolated NGFR positive FOXP3 KO and control Tregs from 10 HD (Fig. 7A). We observed 3 days post TCR and CD28 stimulation an enhanced expansion of FOXP3 KO Treg (Fig 7B). Moreover, we found a slight but significant increase in the proportions of TSDR demethylated cells in FOXP3 KO over control Treg cells (Fig. 7C), which is in line with the increased proliferative capacity in the absence of FOXP3 and confirms high purity of the FOXP3 KO Treg cell. Unfortunately, analyses of CD25 and CD127 markers is not reliable in Treg cell cultures due to IL-2 media supplementation. To assess Teff-like function in FOXP3 KO Treg cell, we collected the supernatant after the 3 days expansion and analyzed production of cytokines including IFN*γ*, TNF*α*, IL-4, IL-5, IL-6, IL-9, IL-10, IL-12, IL-17, IL-18, IL-21, IL-22, IL-23, IL-27 and GM-CSF using the Luminex-multiplexed assay. Cytokine data were normalized to the control:FOXP3 KO cell ratio at the end of the experiment taking into account the higher expansion of the FOXP3 KO Treg. This normalization may favor FOXP3+ Treg cells. We observed increased production of IL-13, IL-17, TNF*α* and GM-CSF by the FOXP3 KO Treg cells (Fig. 7D), suggesting Th2 and Th17 polarization of FOXP3-deficient Treg cells and their potential role in IPEX pathology. Interestingly, we also observed a trend toward higher production of IL-4, which didn’t reach significance because of a possible outlier (Fig. S3). Importantly, we have recently shown that IPEX patients’ T cells are Th2 polarized and that IL-13 and TNF*α* are among the most elevated cytokines in their plasma (Narula M. et al, see references). Collectively, these data indicate that loss of FOXP3 expression in Treg leads to their expansion and production of proinflammatory cytokines, thus providing a possible explanation for expansion of TSDR demethylated cells in IPEX and supporting that loss of Treg identity cells could contribute to the immune dysregulation in IPEX patients.

**Fig. 7.**
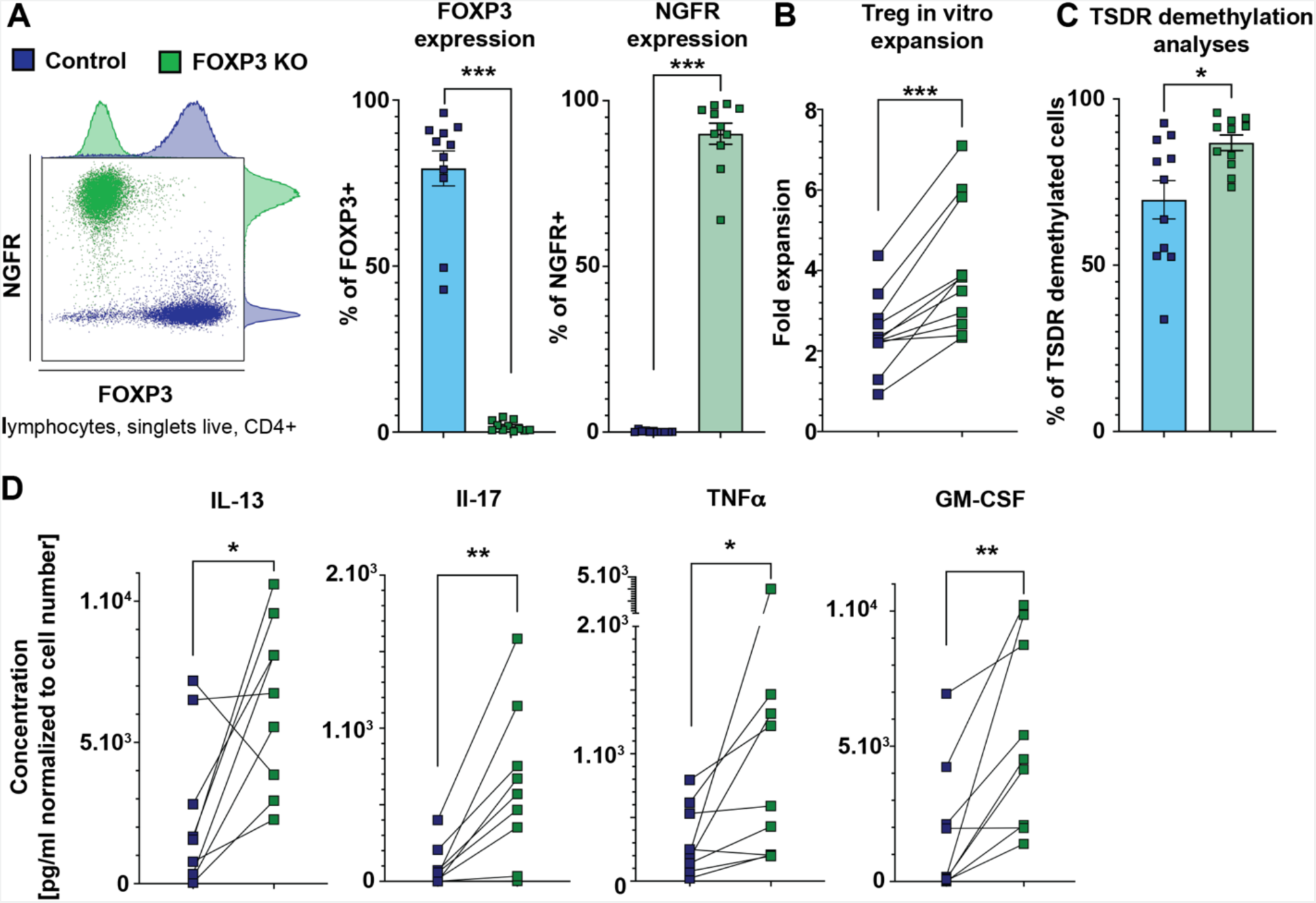
Greater expansion and increased production of proinflammatory cytokines in FOXP3 KO Treg in response to TCR stimulation in vitro. **(A)** Representative FACS plot (left), quantification of FOXP3 (middle), and NGFR (right) expression in FOXP3 deficient and control Treg from 10 HD. **(B)** Treg expansion upon CD3/CD28 stimulation. Cells were counted 3 days post CD3 and CD28 stimulation and the data are plotted as fold increase of cell number from day 0. Lines connect control and FOXP3KO cells from the same donor. **(C)** Quantification of TSDR demethylation analysis in control and FOXP3 deficient Treg. **(D)** Analyses of cytokine production from supernatants of control and FOXP3 KO Treg using a Luminex-based multiplexed assay. Only significantly upregulated or downregulated cytokines are shown. The cytokine concentrations were normalized to the ratio of control:FOXP3 KO Treg cell numbers at the time of supernatant collection for each donor. The data were analyzed using Wilcoxon test (A-D).

## DISCUSSION

In the present work, by using TSDR-demethylation as a method for Treg specific lineage tracing and TCR-repertoire sequencing to evaluate TCR autoreactivity in defined Treg and Teff cell subsets, we show that in patients with IPEX, both FOXP3 deficient Treg and autoreactive Teff are expanded. Moreover, a fraction of the expanded Treg cells lose Treg phenotype and are comprised in the Teff compartment as shown by their Treg-specific epigenetic lineage marker. Collectively, we show that the increased pool of autoreactive T cells has a dual origin, one originating from committed Treg that have lost Treg identity, and the other one, from expansion of autoreactive Teff cells, not appropriately suppressed by Treg cells that are deficient for FOXP3 dependent function.

Despite the normal to low frequency of Treg by standard immunophenotype in IPEX patient (*22–25*), we have recently shown that IPEX patients have increased frequency of TSDR demethylated cells in peripheral blood and that this increase is of great diagnostic importance (*21*)(and Narula M. et al, see references). Our results explain this discrepancy by showing that the Teff compartment of IPEX patients contains TSDR demethylated cells, which are the fraction of IPEX Treg cells that have lost CD25 expression, a crucial cytokine receptor for Treg cell survival. Some of these cells also gain CD127 expression, a key cytokine receptor regulating Teff cell homeostasis (*34, 35*). In addition, we show that some of the Treg cells found in Teff compartment have lost FOXP3 expression, further supporting the concept that a fraction of lineage committed Treg cells in IPEX patients have lost their Treg identity. As a consequence, these cells likely lose regulatory functions. However, they are still derived from the self-Ag specific selected Treg cells, as shown by the TCR analysis and may therefore play a pathogenic role in IPEX. In agreement with this scenario, we observed that self-antigen specific cells were also increased in the Teff2 compartment of IPEX patients, which is composed mainly of TSDR methylated bona fide Teff cells, thus demonstrating expansion of autoreactive bona fide Teff cells in absence of functional Treg cells. Considering that the increase in autoreactivity of Teff2 cell correlates with the loss of Treg identity (TSDR demethylation in Teff1), we speculate that the autoreactive Teff cells expand freely because of the lack of functional Treg cells in patients with FOXP3 mutation (*23*).

A clinically relevant question is, to what extent the loss of Treg identity cells are capable of effector functions. To address this question, we have investigated experimentally whether Treg in the absence of FOXP3 acquire Teff functions, which may be consistent with IPEX patient immune dysregulation. We showed that upon TCR-mediated activation, FOXP3 KO Treg cells upregulate production of the proinflammatory cytokine TNF*α* and Th2 polarizing cytokine IL-13, both of which we found among the most upregulated cytokines in plasma of cohort of IPEX patients with different mutations and clinical manifestations (Narula M. et al, see references). As previously reported, upon TCR stimulation and under neutral conditions, GATA3 is a default response to TCR stimulation in Treg cells (*36*). In addition, GATA3 and FOXP3 could be co-precipitated from the same protein complexes, thus it is plausible to speculate that the dysregulation of IL-13 expression might be a direct GATA3-dependent consequence of FOXP3 loss in the regulation of IL-13 expression (*37*). This finding indirectly suggests that in IPEX patients, *loss of identity Treg cells* are participating in polarization of Teff toward Th2 lineage by secretion of IL-13. As expected, we also observed increased production of IL-17, which reflects the Treg/Th17 imbalance in absence of FOXP3 (*38, 39*). Unlike in Lam et al., we did not observe a broad upregulation of cytokine secretion upon FOXP3 KO in Treg but an increase in specific cytokines (*40*). This includes an upregulation of GM-CSF in FOXP3 KO Treg cells, which has not been reported. GM-CSF production by T cells induce paracrine signaling in phagocytes, their activation, and production of proinflammatory cytokines and chemokines such as IL-1*β*, TNF*α*, CCL17, and CCL22, a mechanism associated with pathogeneses of T cell mediated autoimmune diseases, such as rheumatoid arthritis (RA) and multiple scleroses (MS) (*41–44*). Indeed, in our recent cohort of IPEX patients, we observed increased production of cytokines and chemokines associated with dysregulation of the phagocytic compartment (Narula M. et al, manuscript in revision). Although further work is necessary to fully unravel the role of GM-CSF pathogeneses in IPEX, GM-CSF neutralizing antibodies might be an attractive treatment option, which is currently in clinical trials for RA and MS (*41*). Overall, our in vitro Treg KO data indicates that, upon activation, FOXP3-deficient Treg cells upregulate the expression of effector cytokines and specifically those that we described being increased in IPEX patients. While it is very difficult to clearly establish to what extent autoimmunity is driven by autoreactive Teff cells and to what extent by *loss of identity Treg cells*, these data provide at least a partial explanation for the observed Th2 polarization in IPEX patients and their Th2 and Th17 associated pathologies (ie. eczema and enteropathy, respectively).

Importantly, the increased numbers of Treg cells in IPEX patients are in accordance with the cell intrinsic, inhibitory role of FOXP3 in Treg expansion, as shown by our in vitro experiments. The intrinsic inhibitory role of FOXP3 in Teff cell proliferation has also been reported previously (*45*). Therefore, it is possible that in absence of Treg cells’ suppressive function, the expansion of autoreactive T cells in the periphery may be exacerbated by an intrinsic effect of FOXP3 deficiency in Teff cells.

Interestingly, our discovery of loss of Treg identity cells is partially in line with recently reported single cell RNA sequencing (scRNA seq) data from a different cohort of IPEX CD4^+^ T cells (*46*). The scRNA seq data by D. Zemmour et al. revealed two types of cells with Treg-like transcriptional signatures. The first one named A, clustered separately from IPEX Teff cells and resemble HD Treg cells, and a second one termed B, clustered closer to IPEX Teff cells and thus could represent an intermediate stage of the loss of Treg identity cells we describe here. However, the scRNA seq did not reveal the increased number of Treg, plausibly because of the lack of Treg transcriptional signature of the part of loss of Treg identity cells, indicating that some of the loss of Treg identify cells may profoundly lose their Treg-like transcriptional signature and cluster with the true Teff population rather than with the cluster B.

Our data further shed light into the mechanism of the competitive advantage of wild type (WT) Treg over the mutated Treg cells (*47*). Here we show that IPEX carrier mothers’ Teff cells do not contain loss of identity Treg cells. In addition, IPEX carrier mothers have normal numbers of Treg cells (*29*). Consistently, functional Treg suppress loss of Treg identity and expansion of FOXP3-deficient Treg in the case study of the IPEX patient post transplantation. Therefore, it is likely that FOXP3-deficient Treg cells are being suppressed by functional Treg as they lose the Treg phenotype and become autoreactive Teff-like cells in chimeric patients post transplantation and in carrier mothers.

Our work provides evidence that in the prototypic monogenic autoimmune disease, it is feasible to track Treg cells which lose Treg identity and combines this analysis with TCR receptor sequencing in epigenetically defined Treg and Teff cells subsets. It remains open the question if the loss of Treg identity phenomenon is also present in other more common autoimmune diseases. Data by Ricardo C. Ferreira et all suggest that FOXP3+ cells are abnormally present among the CD25 low T cells in peripheral blood of patients with more common autoimmune diseases(*48*). However, whether these cells can further lose FOXP3 expression and become Teff-like cells, similarly to the IPEX loss of Treg identity cells, remains to be fully determined. Here we showed that similarly to IPEX, the autoreactive Teff cells are also present in APECED, but the mechanisms by which autoreactive clones end up in the Teff compartment are different from IPEX. In IPEX, loss of Treg identity and expansion of autoreactive Teff in the periphery are responsible for the increase in Teff TCR autoreactivity, whereas autoreactive Teff cells in APECED are the result of defective central T cell selection of the Treg/Teff TCR repertoire in the thymus. Therefore, these data are suggesting that the presence of loss of Treg identity cells is a distinctive mechanism of increased autoreactivity in a FOXP3 deficient immune system.

Collectively, these data contribute to the basic knowledge about the role of FOXP3 i) as a gate keeper of Treg identity and ii) in preventing autoreactive Teff cell expansion in humans. In addition, these data will also be of great interest from a translational perspective to monitor impact of novel Treg cell therapies not only in IPEX disease but also in other autoimmune diseases, in which the loss of Treg identity and expansion of autoreactive T cells are still poorly defined.

## MATERIALS AND METHODS

### Study design, Patient and healthy donor PBMC

Patient and healthy control samples were collected upon informed consent on an institutional review board approved protocol (IRB #34131) of the Center for Genetic Immune Diseases, Stanford, CA. Healthy donors were sex matched siblings, parents, and pediatric healthy subjects. In addition, adult healthy donors were obtained on commercial basis from Stanford blood center. For AIRE patient analyses, parents were used as a control if available (4 out of 5 samples, non-sex matched). Patients and healthy donors’ details are summarized in Table 1.

### FACS analyses and sort

Ficoll-Paque™ PLUS media (fisher scientific) isolated peripheral blood mononuclear cells (PBMC) from whole blood, buffy coat or Leukoreduction System Chambers were stained 30 minutes on ice, washed with PBS, analyzed and sorted using BD FACSAria™ III Cell Sorter (BD). Antibodies used in this study are summarized in Table S1. Staining of intracellular antigens was done using Foxp3 / Transcription Factor Staining Buffer Set (eBioscience^TM^) according to manufacturer recommendations. FACS data were analysed using FlowJo software (FlowJo LLC)

### TSDR demethylation analyses

Sorted Teff cell pellets were frozen in PBS. Sorted Treg cells from HD and IPEX patients were stimulated with Dynabeads™ Human T-Activator CD3/CD28 (1:1 beads:Treg ratio, Gibco), and cultured in X-VIVO 15 with Gentamicin, L-Gln, Phenol Red (Lonza), supplemented with 5% human serum (Millipore Sigma) and 300 IU/ml IL-2 (Peprotech) for 5-10 days. Cell pellets were thawed, DNA was isolated, bisulfite converted and analyzed using qPCR as described previously (*49*). Data were analyzed by Epimune GmbH, Germany. Whole blood samples from transplanted patients were normalized to number of CD4^+^ T cells, also determined by epigenetic analysis (*49*).

### TCR analyses

Sorted cell pellets were frozen in PBS. Upon thaw, DNA was isolated using DNeasy Blood and Tissue Kit (Qiagen). TCR CDRB3 region was sequenced by Adaptive biotechnologies. Data were processed by Adaptive and initial analysis was done using immunoSEQ analyzer online tool (Adaptive biotechnologies). The analysis of frequency of autoreactivity promoting and limiting doublets was performed as described previously (*28*). Briefly, TCR CDR3 sequences shorter than 8 amino acids were removed, because of the conserved amino acidic residues at the end of the CDR3 loop. Frequency of autoreactivity promoting and limiting doublets defined by B. D. Stadinski et al., were calculated as a sum of frequencies of productive rearrangements of given set of doublets (*7*). TCR CDRB3 sequences are available online at (data will be available online at Adaptive biotechnology webpage upon manuscript publishing).

### FOXP3 knock out in primary Treg cells, proliferation and cytokine production

To isolate CD4^+^CD25^high^CD127^-^ Treg cells, we used PBMC from LRS chambers and enriched Treg cells using the CD4^+^CD25^+^CD127^dim/-^ Regulatory T Cell Isolation Kit II, human (Miltenyi Biotech) followed by FACS sorting of the CD4^+^CD25^+^CD127^dim/-^ cells to obtain a pure population. Treg cells were stimulated for 7-8 days with Dynabeads™ Human T-Activator CD3/CD28 (1:1 beads:Treg cell ratio, Gibco), and cultured in X-VIVO 15 with Gentamicin, L-Gln, Phenol Red (Lonza), supplemented with 5% human serum (Millipore Sigma) and 300 IU/mL IL-2 (Peprotech). We subsequently edited the cells using Cas9 (Integrated DNA Technologies), chemically modified guide RNA (Synthego corporation), and AAV6 virus (Signagen laboratories, and in house produced (*33*))-mediated delivery of homologous template containing NGFR reporter gene under control of PGK promoter as described previously (*33*). Control Treg cells were treated the same as FOXP3 Knock out Treg cells but were electroporated in absence of CRISPR/Cas9 and non-treated with AAV6. Cells were expanded for 7-10 days and NGFR+ cells were purified using MACSelect™ LNGFR MicroBeads and autoMACS® Pro Separator (Miltenyi Biotech). 2 out of 10 donors were expanded for additional 9 days to obtain sufficient cell number for analyses. Subsequently, 1×10^5^ cells were plated in technical duplicates or triplicates, stimulated with Dynabeads™ Human T-Activator CD3/CD28 (1:1 beads:Treg cell ratio, Gibco), and analyzed for cell count, FOXP3 expression, and cytokine production 3 days post stimulation. Samples with very low TSDR demethylation were removed from analyses.

### Cytokine production analyses

Supernatants were collected 3 days post Dynabeads™ Human T-Activator CD3/CD28 (1:1 beads:Treg cell ratio, Gibco) stimulation for analysis of cytokine production using a Luminex-based multiplexed assay (Th1/2/9/17 18-plex Human ProcartaPlex Panel, ThermoFisher) per manufacturer recommendation. Technical duplicates were averaged and for each donor the values were normalized to control:FOXP3 KO cell number ratio at the time of supernatant collection 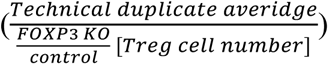.

### Statistical analyses

Statistical analyses were performed using GraphPad prism software version 9 (GraphPad Software, Inc.). When samples were analyzed in technical duplicates or triplicates, only averages were plotted and used for analyses. Statistical tests used in this study are always indicated in the figure legend. Significance is indicated for p value p<0.05 = *, p<0.01 = **, p<0.005 = ***, p<0.0001 = ****. N represents number of donors or values per group.

## Supplementary Materials

Fig. S1. FOXP3 expression in patients analyzed in Fig.1E

Fig. S2. TCR beta gene usage in IPEX and HD Treg, Teff1 and Teff2 populations.

Fig. S3. Cytokine production by FOXP3 KO and control Treg cells.

Table S1. List of antibodies.

**Narula M.** *et.al.*, Epigenetic and Immunological Indicators of IPEX Disease in subjects with FOXP3 gene mutation. Manuscript under revision.

## Acknowledgments

We would like to thank the patients and families, who made this study possible by kindly donating their blood. In addition, we would like to thank to Robert Artur Freeborn and Benjamin Thomas for English corrections, Laura Passerini for carrier mother’s sample collection, Kenneth Weinberg for providing us Aria II instrument, and Catherine Carswell-Crumpton and Cheng Pan from FACS core at Institute for Stem Cell Biology and Regenerative Medicine for kind technical support.

## Funding

This study was supported by:

Maternal and Child Research Institute (MCHRI) grant: New idea to RB.

Maternal and Child Research Institute (MCHRI) grant Postdoctoral support to SB.

Bonnie Uytengsu and family endowment for the Center for Genetic Immune Diseases (CGID) and the Center for Definitive and Durative Medicine (CDCM).

## Author contributions

Conceptualization: SB, RB, EM, EL

Methodology: SB, RB, EM, EL, MN

Investigation, sample collection, processing and data analyzes: SB, EL, UL, MM, MN, AR, JB, JS, SO, LM, YG, MB, AB, RB.

Visualization: SB, RB

Funding acquisition: RB, SB, MGR

Supervision: RB, EM

Writing – original draft: SB, RB, EM, MM

Writing – review & editing: SB, RB, EM, MM, AB, EL, MN, JS, SO

## Data and materials availability

All major data are available in the main text or the supplementary materials. Additional data are available upon request from the corresponding author.

