## Supplementary material for "Loss of FOXP3 function causes expansion of two pools of autoreactive T cells in patients with IPEX syndrome": S_Fig.pdf

**Fig. S1. FOXP3 expression in patients analyzed in Fig.1E.** Treg, Teff1, and Teff2 cells were gated as in Fig. 1A.

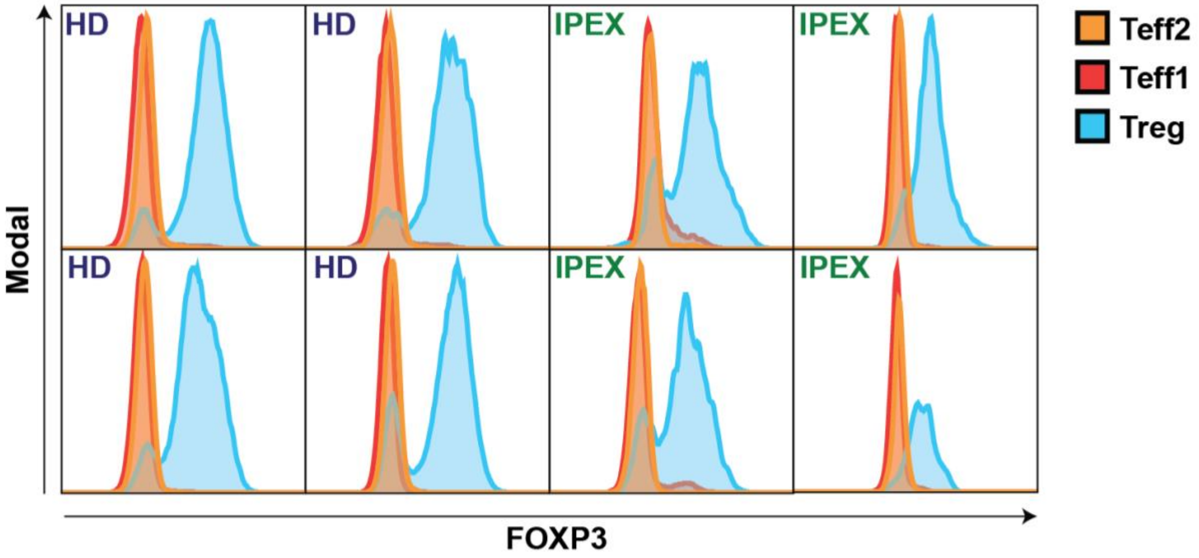

**Fig. S2. TCR beta gene usage in IPEX and HD Treg, Teff1 and Teff2 populations.**

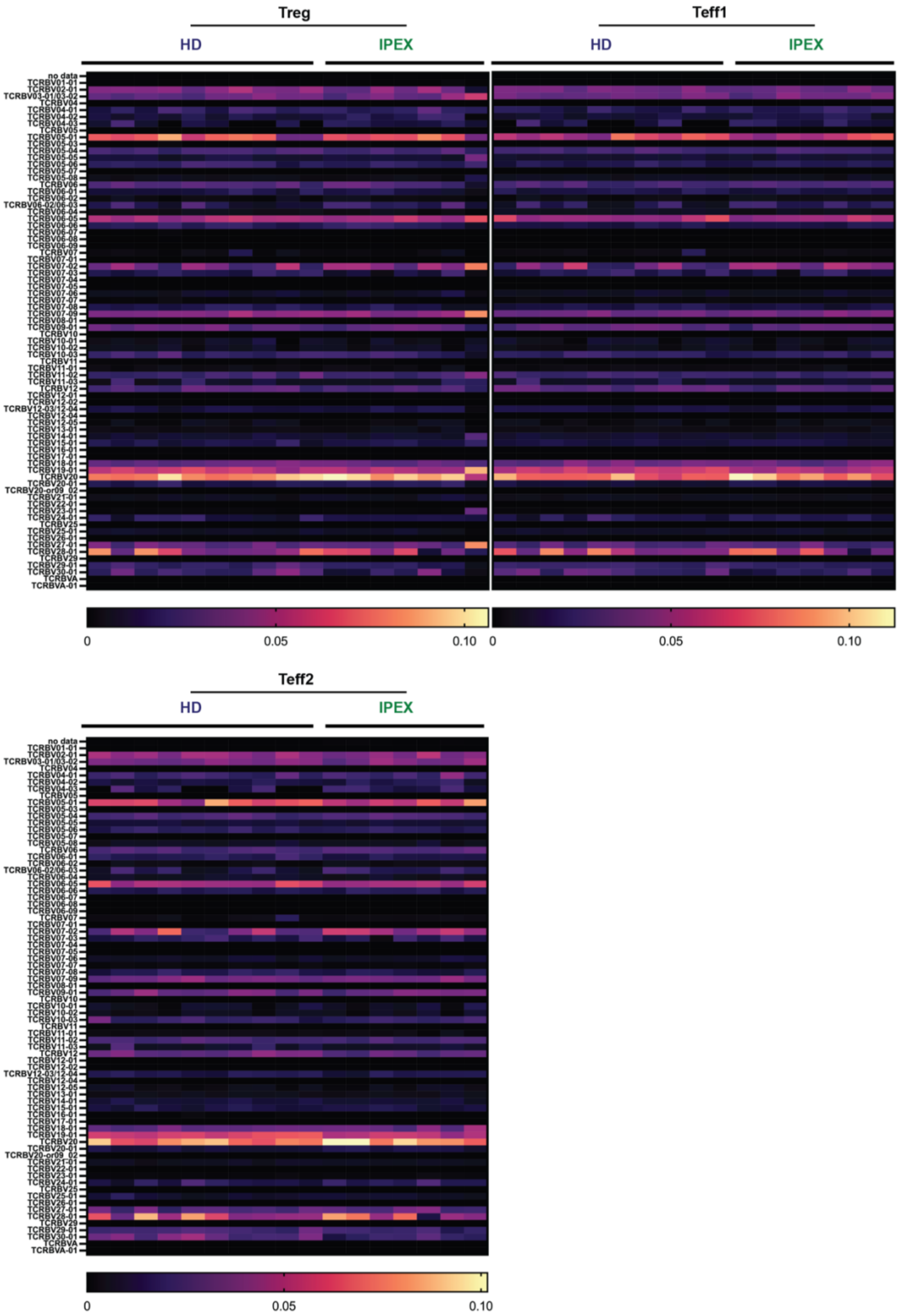

**Fig. S3. Cytokine production by FOXP3 KO and control Treg cells.**

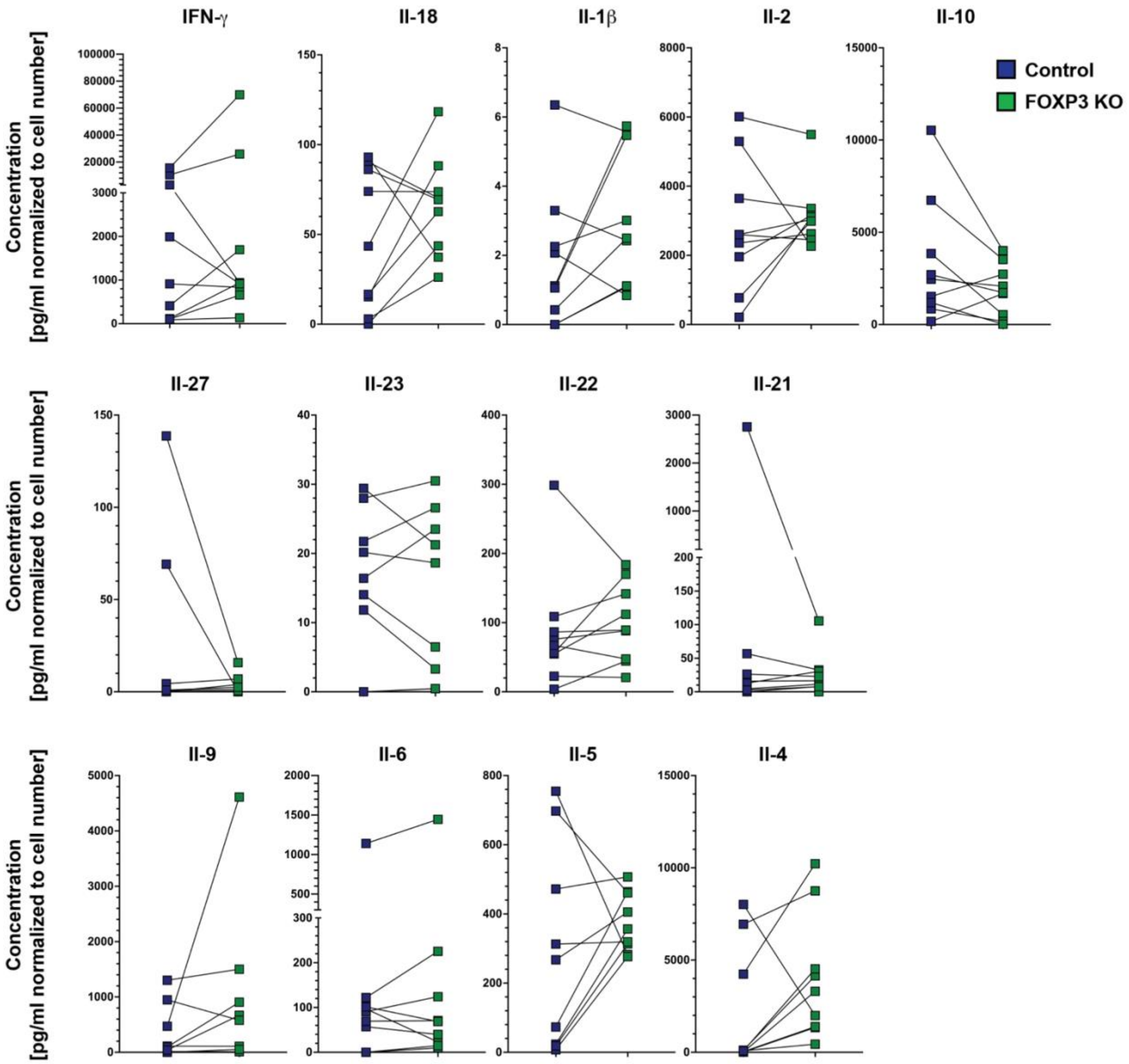

**Table S1. List of antibodies**

| Antigen/reagent | Clone/details | Conjugate | Company |
|---|---|---|---|
| CD3 | OKT3 | PE,BB700 | biolegend, BD |
| CD4 | RPA-T4 | a700 | BD |
| CD14 | 63D3 | PB | biolegend |
| CD127 | A019D5 | PE/Cy7 | biolegend |
| CD25 | 2A3 | APC, FITC, BV605 | BD |
| FOXP3 | 259D, 150D | a488, PE | biolegend |
| NGFR | C40-1457 | PE-CF594 | Biolegend |
| HLA-A2 | BB7.2 | APC | Biolegend |
| Live/Dead Fixable Aqua | Cell Stain Kit | | ebiosciences |
| Hoechst 33258 | Staining Dye Solution (ab228550) | | Abcam |
| LIVE/DEAD™ Fixable Violet | Cell Stain Kit | | ebiosciences |
